# Chemotaxis and selective interactions of *Trichomonas vaginalis* with the vaginal bacteria

**DOI:** 10.64898/2026.03.25.714215

**Authors:** Manuela Blasco Pedreros, Maria Florencia Irigoyen, Augusto Simoes-Barbosa, Angelica Montenegro Riestra, Natalia de Miguel

## Abstract

*Trichomonas vaginalis* is an extracellular parasite that inhabits the human genital tract, yet little is known about how it senses and responds to the complex vaginal microbial ecosystem. Here, we show that *T. vaginalis* exhibits chemotactic behavior on semisolid surfaces, forming multicellular assemblies that coordinate collective migration. Parasite colonies display both positive and negative chemotactic responses, indicating the ability to detect and react to diffusible signals. Different parasite strains display marked mutual avoidance between neighboring colonies, highlighting specific recognition mechanisms. Furthermore, we show that *T. vaginalis* is strongly attracted to acidic environments, revealing a niche-adapted pH taxis. Given that vaginal bacteria critically shape local pH, we examined parasite responses to representative members of the vaginal microbiota. *T. vaginalis* exhibited preferential chemotactic migration toward *Lactobacillus gasseri*, a hallmark species of eubiotic community state types (CSTs), over *Gardnerella vaginalis*, which is associated with dysbiotic CST-IV communities, while showing no detectable attraction to *Escherichia coli*. This selective migration correlated with a robust chemotactic response to lactic acid, a major metabolite produced by lactobacilli. Additionally, when the parasite is co-cultured with the equal number of *L. gasseri* and *G. vaginalis*, *T. vaginalis* exhibits a clear preferential binding to *L. gasseri*, as demonstrated by flow cytometry and fluorescent microscopy. We show that co-culture of *T. vaginalis* with either *L. gasseri* or *G. vaginalis* results in enhanced parasite growth only in the presence of *L. gasseri*. Collectively, these findings reveal pH taxis; bacteria-directed migration and preferential association with Lactobacillus as previously underappreciated behavioral traits of *T. vaginalis*. Such behaviors may destabilize protective microbial communities and drive the transition toward a CST-IV–type dysbiotic state which is frequently associated with trichomoniasis.

## Introduction

Microorganisms behave as social entities in their natural enviroments, and are capable of communicating with one another while engaging in cooperative behavior (Bassler & Losick, 2006; Shapiro, 1998). Well-characterized social activities include biofilm formation, social motility, fruiting body development and quorum sensing (Bassler & Losick, 2006; Harshey, 2003; Kaiser, 2003; Shapiro, 1998), which confer specialized functionalities and offer advantages over a unicellular lifestyle. Some of these advantages include increased protection from external antagonists, such as desiccation or host defenses, access to nutrients, exchange of genetic information and enhanced ability to colonize, penetrate and migrate on surfaces (Bassler & Losick, 2006; Harshey, 2003; Shapiro, 1998). Social interactions in bacterial and fungal pathogens, strongly influence microbial physiology and pathogenesis, and the recognition of social behavior as a widespread bacteria feature has profoundly changed our understanding of microbiology (Bassler & Losick, 2006; Kaiser, 2003; Shapiro, 1998). Specifically, surface-induced collective motility, often termed social motility (SoMo) have been implicated in a wide range of biological functions in biofilm-forming bacteria (e.g., Pseudomonas and Vibrio spp.), in fruiting-body forming bacteria (e.g., Myxococcus) or in eukaryotes such as Dictyostelium (Bastin & Rotureau, 2015; Fraser & Hughes, 1999; Harshey, 2003; Kaiser, 2003; Kohler et al., 2000). However, in parasitic protozoa such behaviors have only been reported in *Trypanosoma brucei*, where SoMo has been linked to environmental sensing and pH taxis (Bastin & Rotureau, 2015; Oberholzer et al., 2010a; Reuner et al., 1997). Whether other species of pathogenic protozoa exhibit analogous forms of collective migration remains unknown.

*Trichomonas vaginalis* is an extracellular protozoan parasite that causes the most common non-viral sexually transmitted disease (WHO, 2021). It colonizes the mucosal surfaces of the human genitourinary tract. Although asymptomatic infection is common, multiple symptoms and pathologies can arise in both men and women, including vaginitis, urethritis, prostatitis, premature rupture of membranes and preterm delivery of low-weight infants and infertility (Fichorova, 2009; Swygard et al., 2004). Moreover, *T. vaginalis* has been identified as a key cofactor in enhancing HIV transmission, as individuals infected with the parasite exhibit a significantly higher risk of acquiring and transmitting the virus (McClelland et al., 2007; Van Der Pol et al., 2008). Chronic *T. vaginalis* infection has also been associated with increased risk of cervical and aggressive prostate cancers (Gander et al., 2009; Twu et al., 2014). While it is well established that *T. vaginalis* interacts extensively with its host to promote survival and transmission, recent studies have begun to reveal that communication among parasites themselves also plays an important role (Kochanowsky et al., 2024; Salas et al., 2023). Different *T. vaginalis* strains can exchange information through extracellular vesicles and cytoneme-like membranous connections (Kochanowsky et al., 2024; Salas et al., 2023). Communication between isolates with distinct phenotypic traits has been shown to alter the adherence capacity of recipient parasites, potentially leading to phenotypic shifts with clinical implications (Salas et al., 2023; Twu et al., 2013). Despite growing recognition of cell-to-cell communication in *T. vaginalis* pathogenesis, communication and sensing at the population level, as well as with the surrounding vaginal bacteria, have yet to be explored.

The vaginal tract is a dynamic environment that varies with the hormonal status across ages, featuring high glycogen and mucin content, and a low pH in healthy women of reproductive age (Landolt et al., 2025; Zhai et al., 2025). The vagina typically carries a bacterial load of approximately 10⁷–10⁸ CFU/g of vaginal fluid (Pacha-Herrera et al., 2020; Phukan et al., 2018a). While the dominance of a single species of *Lactobacillus* is frequently observed; a greater microbial diversity, encompassing both commensal and potentially pathogenic bacteria has also been described in the vaginal microbiota of these women (Amabebe & Anumba, 2018a). This microbiota has been classified into five community state types (CSTs). About 75% of asymptomatic women of reproductive age harbor a CST that is dominated by a single *Lactobacillus* species: *Lactobacillus crispatus* (CST-I)*, L. gasseri* (CST-II)*, L. iners* (CST-III)*, or L. jensenii* (CST-V) (Amabebe & Anumba, 2018a; Ravel et al., 2011). Among these *Lactobacillus*-dominated CSTs, *L. gasseri* is considered the most stable over time, rarely transitioning to other community types (Ravel et al., 2011). The fifth community (CST-IV), present in approximately 25% of these women, lacks a dominant *Lactobacillus* species and is instead characterized by a diverse consortium of mostly anaerobic bacteria, including *Prevotella bivia, Atopobium vaginae,* and *Gardnerella vaginalis* (Z. (Sam) Ma & Li, 2017; Ravel et al., 2011). The depletion of protective *Lactobacillus* species and the overgrowth of these anaerobes is a hallmark of bacterial vaginosis (BV), a dysbiotic polymicrobial condition that has significant consequences to reproductive health. Notably, trichomoniasis is commonly associated with this CST-IV microbiome suggesting a relationship between *T. vaginalis* infection and shifts in vaginal microbial community composition (Dong et al., 2024). Despite this association, the directionality of this relationship remains unclear; that is whether CST-IV composition facilitates the parasite infection, or whether *T. vaginalis* itself drives the vaginal community toward a CST-IV state. Regardless of these possibilities, we envisage that the parasite may sense bacterial-derived cues and actively respond to these at a population level in order to initiate bacterium-specific interactions. To our knowledge, this has not been investigated yet. In this study, we investigate whether *T. vaginalis* exhibits directed chemotaxis and selective interactions with vaginal bacteria, providing new insight into parasite–microbiota crosstalk.

To fill this gap of knowledge, here we investigated for the first time whether *T. vaginalis* exhibits directed chemotaxis and selective interactions with members of the vaginal microbiota. Using a semi-solid surface model, we examine parasite migration behavior and its responses to bacterial and chemical cues representative of the vaginal environment. Our findings reveal that *T. vaginalis* displays collective surface migration and responds to bacterial-associated signals, including acidic metabolites, and exhibits interactions with specific bacterial species. Together, these results provide new insight into parasite:microbiota crosstalk and highlight the potential role of environmental sensing in shaping parasite behavior within the vaginal niche.

## Results

### Collective migration and inter-parasite avoidance in *T. vaginalis*

When cultured on semisolid agarose, *Trichomonas vaginalis* exhibited collective motility, reminiscent of the social motility (SoMo) behavior previously described for *Trypanosoma brucei* (Oberholzer et al., 2010b). During this process, the parasites formed organized multicellular communities that radiated outward from the inoculation site (Fig. 1). In both analyzed strains (NYH209 and B7RC2), parasites at the colony periphery displayed polarized and synchronized movement, consistent with intercellular communication and coordinated population behavior (Fig. 1A and 1C). NYH209 frequently formed coordinated peripheral projections extending around the colony (Fig. 1A), whereas B7RC2 typically expanded as a compact colony without the presence of prominent projections (Fig. 1C). When two expanding colonies of the same strain were cultured on the same plate and approached each other, their advancing fronts frequently halted or changed direction, indicating active avoidance (Fig. 1B and 1D). Parasites within these projections actively changed direction or stopped migrating upon sensing nearby *T. vaginalis* populations (Fig. 1B and 1D). This effect is observed with both analyzed strains, NYH209 (Fig. 1B) and B7RC2 (Fig. 1D). The ability of colonies to detect and avoid one another over a distance suggests that *T. vaginalis* responds to diffusible chemical cues released by neighboring groups. A similar avoidance response was observed when colonies from different parasite strains were cultured in proximity (Fig. 2). When parasites from NYH209 strain were confronted with B7RC2, CDC1132, or B7268 strains, the migrating fronts altered their trajectories, indicative of negative chemotaxis between distinct *T. vaginalis* populations (Fig. 2). As observed, NYH209 colonies halted or changed direction upon approaching colonies of different strains, indicating that active avoidance is not strain-dependent (Fig. 2). Together, these observations demonstrate that *T. vaginalis* exhibits negative chemotaxis during social motility, adjusting its migration to prevent intercolony contact.

**Figure 1.**
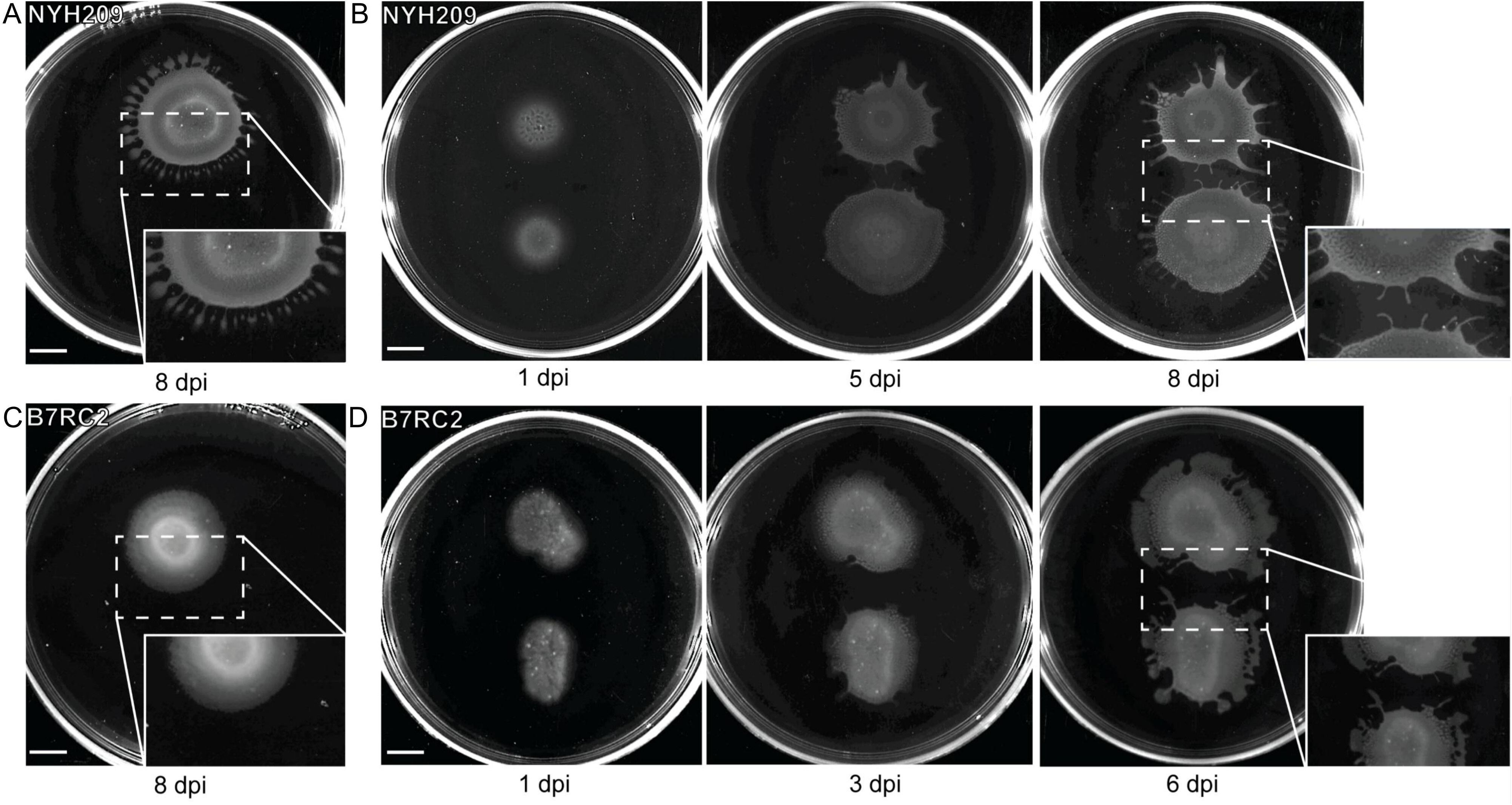
Social motility in *T. vaginalis.* (A) Representative images of colony expansion of the NYH209 strain on semi-solid agarose plates, monitored for up to 8 days post-inoculation (dpi). Insets show higher-magnification views of the leading-edge morphology and multicellular organization. (B) Two NYH209 colonies plated in close proximity and monitored for 8 dpi. Insets highlight the redirection of the leading edge as colonies approach one another at later time points. (C) Representative images of colony expansion of the B7RC2 strain on semi-solid agarose plates, monitored for up to 8 dpi. Insets emphasize differences in leading-edge morphology. (D) Two B7RC2 colonies plated in proximity and followed for 8 dpi. Insets show leading-edge redirection upon intercolony approach at later stages. Images are representative of four independent experiments. Scale bar, 10 mm.

**Figure 2.**
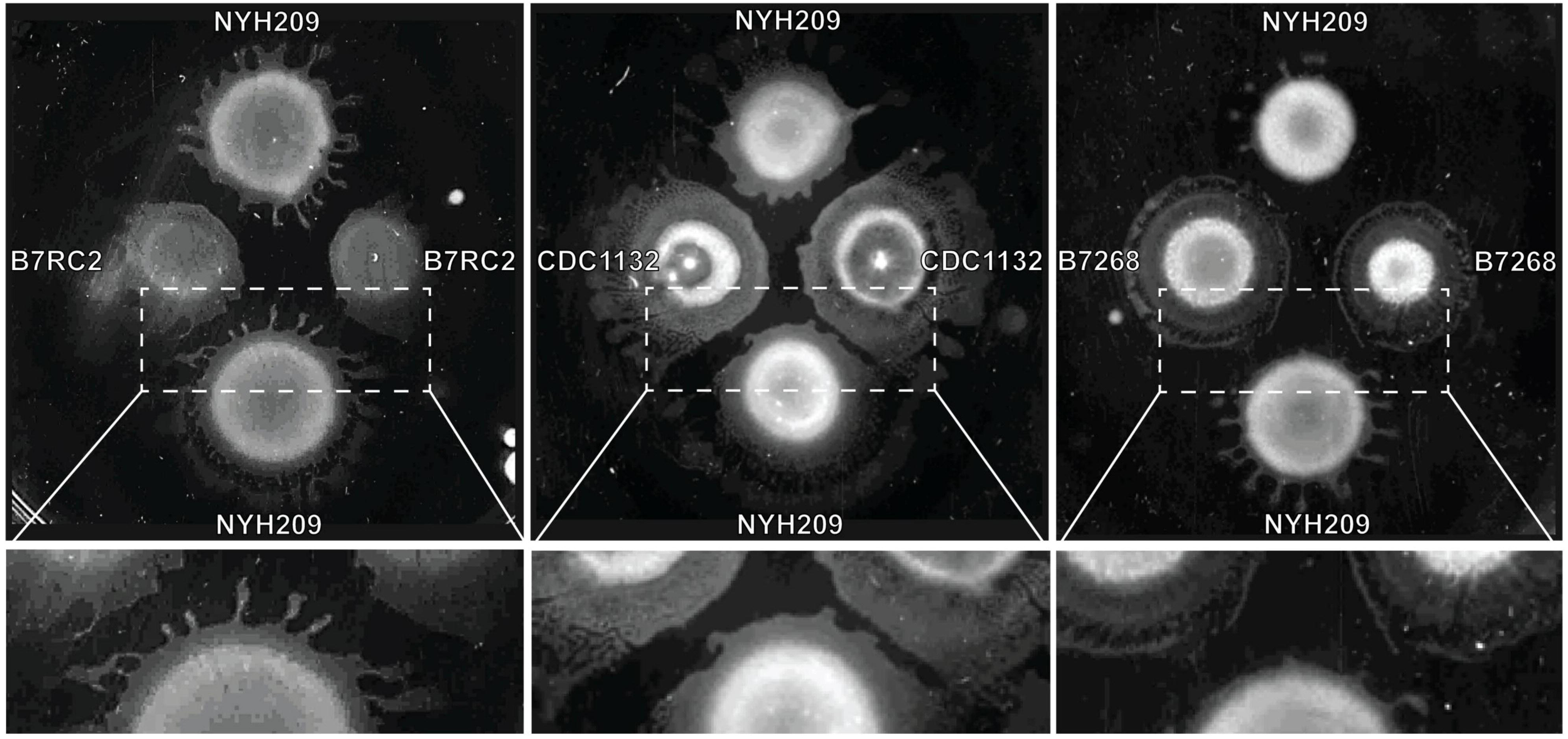
Strain-specific interactions during social motility in *Trichomonas vaginalis*. Representative images of semi-solid agarose plates showing social motility patterns of different *T. vaginalis* strains plated in proximity and monitored over time. The NYH209 strain was co-cultured with B7RC2 (left panel), CDC1132 (middle panel), or B7268 (right panel). Colony expansion and directional growth were assessed by examining changes in the leading-edge morphology as colonies approached one another. Representative images show NYH209 colonies confronted with B7RC2, CDC1132, or B7268 strains at 6 days post-inoculation (dpi). Dashed boxes indicate regions of interaction between expanding colonies, and lower panels show higher-magnification views of these areas. Images are representative of three biologically independent experiments.

### *T. vaginalis* exhibits pH taxis

As it has been recently proposed that the social motility (SoMo) behavior of *Trypanosoma brucei* on semisolid surfaces is driven by pH taxis (Shaw & Roditi, 2023), we investigated whether *T. vaginalis* migration is influenced by pH gradients. To test this, strains NYH209 and B7RC2 were inoculated onto semisolid plates, and evaluated if manipulating the pH gradient had an influence on migration (Fig 3A and B). To this end, 3 × 10^6^ parasites were inoculated onto plates and solutions of 1M HCl or NaOH were pipetted onto the surface at a 2 cm distance (Fig. 3A and B). The plates were incubated for 6 days. As can be observed in Fig. 3, HCl acted as a chemoattractant, causing projections to reorient toward the acid, while NaOH triggered avoidance or reversal of migration (Fig 3). These results demonstrate that *T. vaginalis* exhibit pH taxis, with acid acting as an attractant. This behavior is consistent with the acidic vaginal environment where the parasite naturally resides and contrasts with *T. brucei*, which displays migration toward more alkaline conditions (Shaw et al., 2022; Shaw & Roditi, 2023).

**Figure 3.**
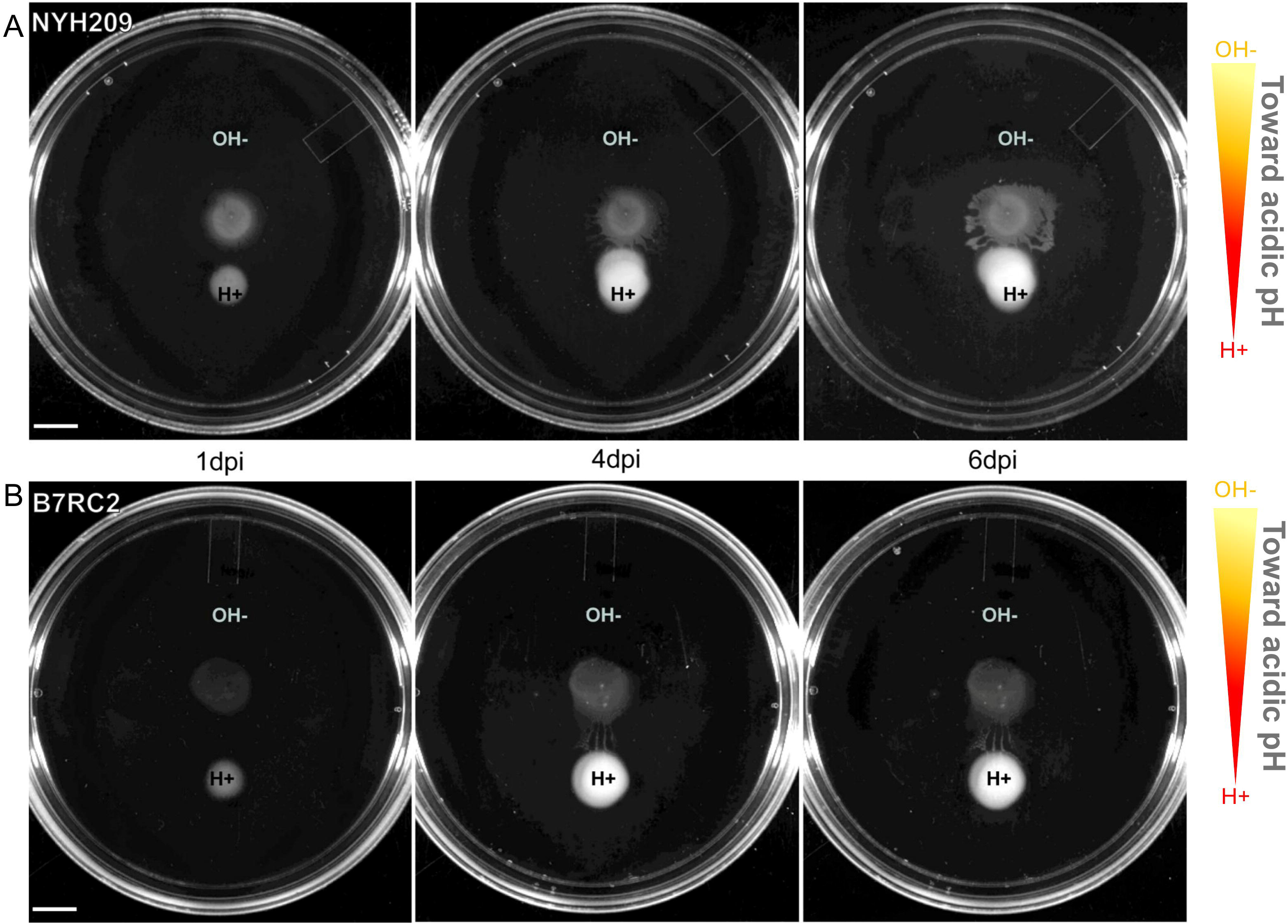
*T. vaginalis* exhibits pH-directed migration on semi-solid agarose. Parasites were inoculated onto semi-solid agarose plates and exposed to acidic and alkaline cues. Parasites (3 × 10⁶ cells) were seeded at the center of the plate, and HCl (H^+^) or NaOH (OH^-^) solutions were applied onto the agarose surface at a 2 cm distance. Representative images of the NYH209 (A) and B7RC2 (B) strains at 1, 4 and 6 days post-inoculation (dpi). Images are representative of three independent experiments. Scale bar, 10 mm.

### *T. vaginalis* exhibit positive chemotactic responses toward vaginal bacteria

Trichomoniasis is typically associated with CST-IV, a dysbiotic microbiome characterized by the depletion of host-protective commensals lactobacilli and the expansion of anaerobic bacterial consortium that is comparable to the one seen for bacterial vaginosis (Artuyants et al., 2024; Cardoso & Tasca, 2025; Hinderfeld et al., 2019). Given that *T. vaginalis* coexists with the host vaginal microbiome, we evaluated its social motility behavior when encountering bacterial colonies commonly found in the vagina. Three representative species of bacteria were specifically chosen for this study: 1. *Lactobacillus gasseri* (CST II, a hallmark of vaginal eubiosis that rarely transition to other CSTs (Gajer et al., 2012); 2. *Gardnerella vaginalis* (CST IV, a key species in vaginal dysbiosis, a primary pathogen associated with BV (Gilbert et al., 2013; Onderdonk et al., 2016) which often accompanies women with trichomoniasis (Brotman et al., 2012; Ravel et al., 2011) and 3; *Escherichia coli* (bacterium commonly associated with urinary tract infections). When *T. vaginalis* colonies from NYH209 and B7RC2 strains were exposed to these bacteria on agar plates, parasite projections actively migrated toward the *L. gasseri* and *G. vaginalis* bacterial colonies, frequently altering their trajectories to make direct contact (Fig 4). These results indicate that parasite movement toward bacteria reflects an active chemotactic response rather than a passive lack of avoidance. Interestingly, at early incubation times, both analyzed strains exhibited significant stronger chemotactic response toward *L. gasseri* than *G. vaginalis*, suggesting a preferential attraction (Fig 4B and 4D). After longer incubation periods, this difference became less apparent as chemotaxis toward both *L. gasseri* and *G. vaginalis* was observed in both analyzed strains (Fig. 4B and 4D). In contrast, migration is not observed toward *E. coli* at any analyzed time points, supporting selective sensing of vaginal bacteria rather than a generalized response to bacterial cues (Fig 4A and 4C).

**Figure 4.**
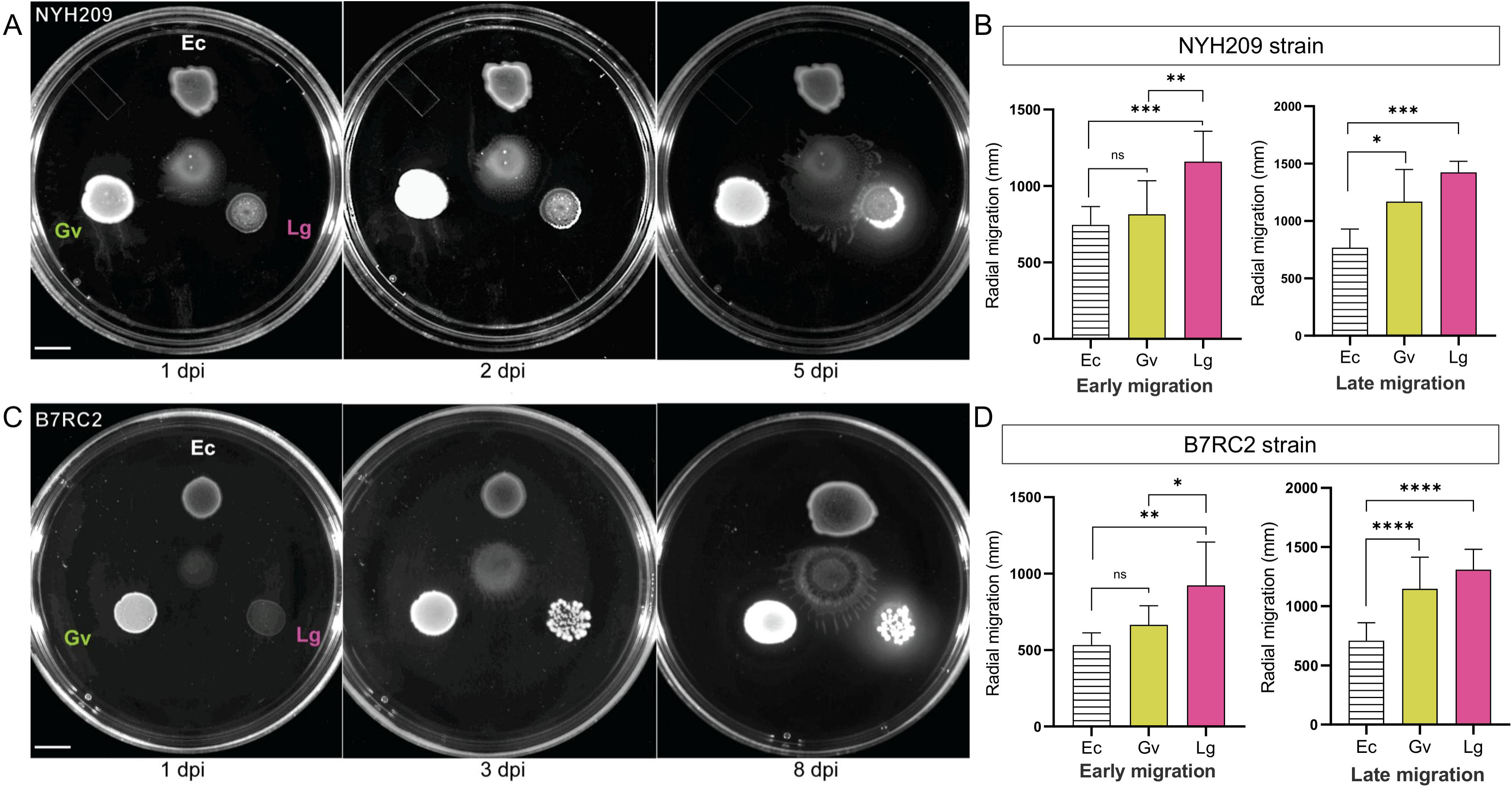
*T. vaginalis* displays selective chemotactic migration toward vaginal bacterial species. (A) Representative images of NYH209 parasite colonies exposed to *E. coli* (Ec), *G. vaginalis* (Gv), or *L. gasseri* (Lg) at 1, 2, and 5 days post-inoculation (dpi). (B) Quantification of radial migration of NYH209 during early (2 dpi) and late (5 dpi) migration. (C) Representative images of B7RC2 colonies exposed to *E. coli* (Ec), *G. vaginalis* (Gv), or *L. gasseri* (Lg) at 1, 3, and 8 dpi. (D) Quantification of radial migration of B7RC2 during early (3 dpi) and late (8 dpi) migration. Quantitative data are presented as mean ± SD. Statistical significance was determined using ordinary one-way ANOVA followed by multiple comparisons tests. Significance levels are indicated as P < 0.05 (*), < 0.01 (**), < 0.001 (***), and < 0.0001 (****); ns, not significant. Images are representative of three independent experiments. Scale bar, 10 mm.

Considering the negative chemotaxis toward *E. coli* (Gram-negative) and the positive response toward Gram-positive bacteria, we evaluated whether *T. vaginalis* sensing and attraction are restricted to Gram-positive bacteria present in its natural niche. To this end we compared the parasite social motility behavior in the presence of *Lactobacillus gasseri* (Gram-positive), *Gardnerella vaginalis* (Gram-variable), and *Bacillus methylotrophicus* (Mt3), a Gram-positive bacterium commonly used in agricultural applications (Romero et al., 2016; Vignatti et al., 2019). As can be observed in Figure 5, colonies of NYH209 and B7RC2 strains exhibited positive chemotaxis toward all three bacterial species (Fig. 5), indicating that attraction is not limited to bacteria present in the vaginal environment.

**Figure 5.**
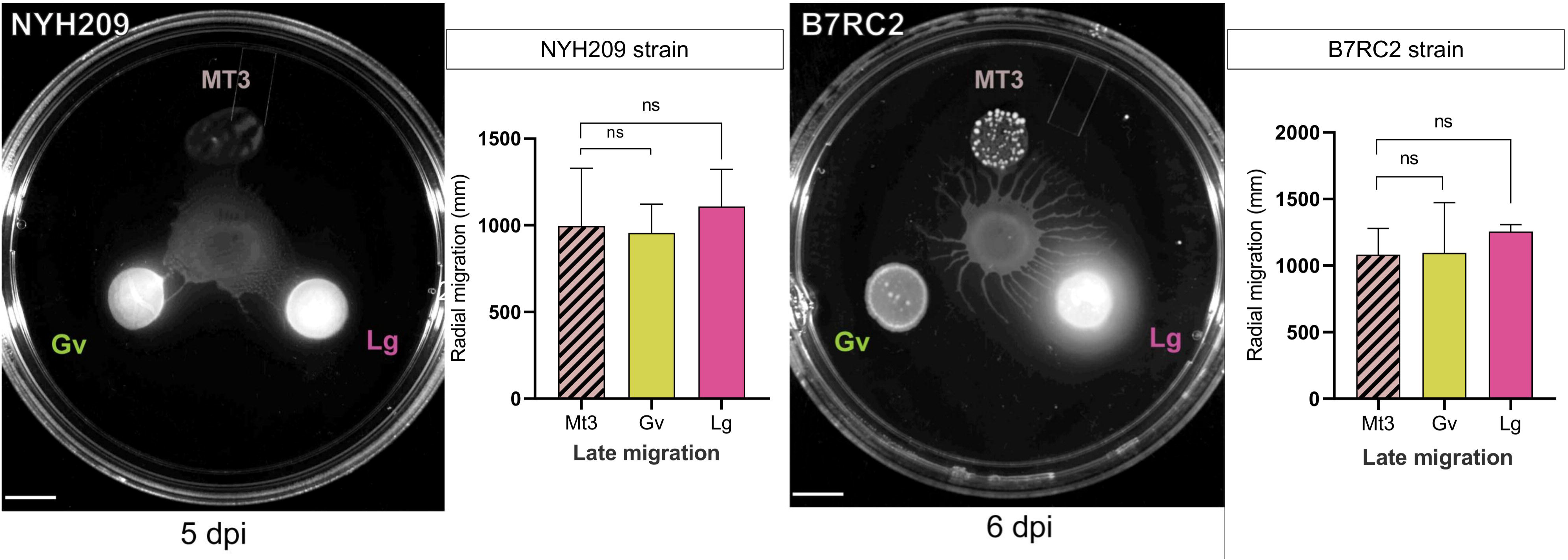
Directional migration of *Trichomonas vaginalis* toward Gram positive bacterial species during social motility. Images of semi-solid agarose plates showing directional migration of *T. vaginalis* strains NYH209 (left) and B7RC2 (right) in the presence of three bacterial species positioned at fixed peripheral locations: *Bacillus methylotrophicus* (MT3), *G. vaginalis* (Gv), or *L. gasseri* (Lg). Parasites were inoculated at the center of the plate and monitored over time (5 dpi for NYH209; 6 dpi for B7RC2). Bar graphs depict quantification of late migration responses of NYH209 and B7RC2 toward each bacterial species. Images are representative of three independent biological experiments. Scale bar, 10 mm.

### Vaginal bacteria modify pH

The vaginal microenvironment is typically dominated by a limited number of Lactobacillus species that maintain an acidic pH of approximately 3.5 - 4.5, thereby creating conditions that are considered protective against pathogen colonization and the development of vaginal dysbiosis (Vaneechoutte, 2017b). Increasing evidence suggests that protection is primarily mediated by lactic acid produced by specific vaginal lactobacilli, rather than by organic acids in general, highlighting a central role for lactobacillus-derived lactic acid in maintaining vaginal health (Tachedjian et al., 2017; Vaneechoutte, 2017a, 2017b). In bacterial vaginosis (BV), this lactobacilli-dominated ecosystem is disrupted and replaced by a diverse consortium of anaerobic bacteria (Cauci et al., 2005; Foxman et al., 2014; Srinivasan et al., 2015). This shift is associated with reduced lactic acid levels and increased concentrations of short-chain fatty acids, such as acetic acid, butyric acid, propionate, and succinate, produced by anaerobes (Al-Mushrif et al., 2000; Amabebe & Anumba, 2018b; Chaudry et al., 2004). Considering the observed attraction of *T. vaginalis* to *L. gasseri* and *G. vaginalis* colonies, as well as its preference for acidic pH, we first assessed the ability of each bacterial species to acidify the surrounding environment. To this end, *L. gasseri*, *G. vaginalis*, and *E. coli* were cultured for 48 hours, and the pH of the medium was monitored over time. Both *L. gasseri* and *G. vaginalis* acidified their environment *in vitro* more than *E. coli* (Fig. 6A). Then, we evaluated the chemotactic behavior of the NYH209 and B7RC2 strains in response to lactic acid, acetic acid, or water as a control. Attraction to both acids was observed at 5 days post-inoculation for NYH209 (Fig. 6B) and 4 days post-inoculation for B7RC2 (Fig. 6C). Our findings support a model in which the parasite senses and migrates toward acidic microenvironments generated by vaginal bacteria.

**Figure 6.**
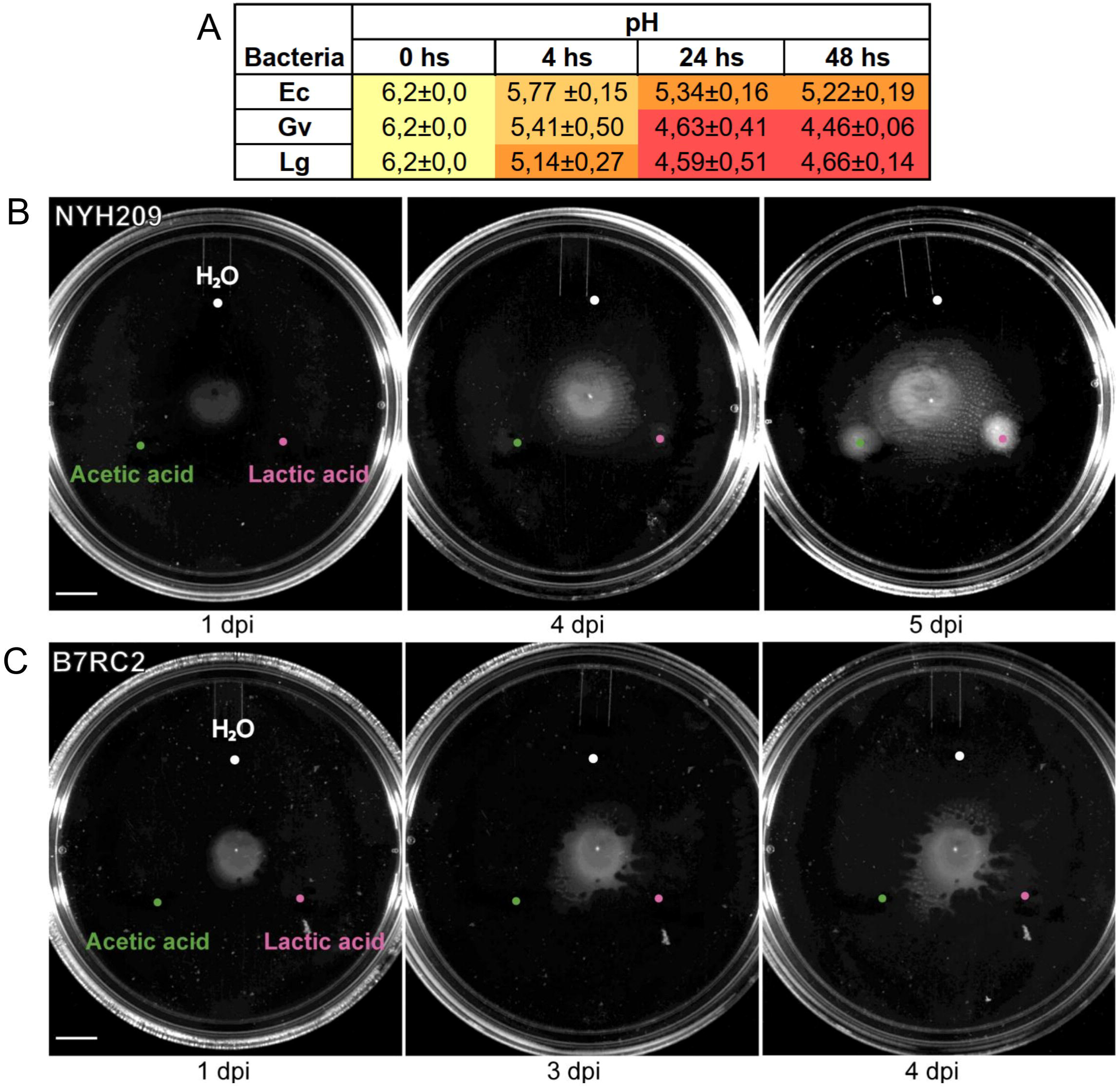
Lactic and acetic acid cues act as chemoattractants for *T. vaginalis*, promoting directional migration. (A) pH measurements of bacterial cultures over time (0, 4, 24, and 48 hours). *Escherichia coli* (Ec), *Gardnerella vaginalis* (Gv), and *Lactobacillus gasseri* (Lg). Values represent mean ± SD. (B) Representative images of NYH209 colonies exposed to water (H_2_O), lactic acid, or acetic acid at 1, 4, and 5 days post-inoculation (dpi). (C) Representative images of B7RC2 colonies under the same experimental conditions at 1, 3, and 4 dpi. Parasites (3 × 10⁶ cells) were inoculated at the center of semi-solid agarose plates, and acidic cues were applied onto the agarose surface at 2 cm of the inoculation site. Images are representative of three independent biological experiments. Scale bar, 10 mm.

### *T. vaginalis* preferentially associates with *L. gasseri* over *G. vaginalis*

Given the preferential migration toward *L. gasseri*, we next asked whether *T. vaginalis* exhibits selective interactions with specific members of the vaginal microbiota. To address this, we assessed parasite:bacteria binding using flow cytometry and fluorescence microscopy (Fig. 7A). *L. gasseri and G. vaginalis* were independently labelled with CellTrace Far Red (red) and CFSE (green), respectively, mixed at equal proportions, and co-incubated with *T. vaginalis* (B7RC2 strain) (Fig. 7). Interestingly, when co-cultured with equivalent bacterial numbers, *T. vaginalis* displayed a noticeable preferential association with *L. gasseri*, as demonstrated by both flow cytometry (Fig. 7B-C) and fluorescence microscopy (Fig. 7D-E). As can be observed, the flow cytometry analysis showed a clear and progressive increase in parasite-associated fluorescence derived from the binding of *L. gasseri* (Fig. 7B). Specifically, a significant rise in median fluorescence intensity (MFI) relative to time 0 was detected from 30 minutes post-incubation. In contrast, fluorescence associated with *G. vaginalis* remained low and did not show a comparable increase throughout the time course (Fig. 7B). Quantification of Δmedian MFI confirmed a significantly stronger association of *T. vaginalis* with *L. gasseri* compared to *G. vaginalis* at analyzed time points (Fig. 7C). These findings were confirmed by fluorescence microscopy, which revealed prominent accumulation of *L. gasseri* bound to *T. vaginalis* cells as early as 30 min post-incubation, with more extensive bacterial clustering evident at 120 min (Fig. 7D). In contrast, *G. vaginalis* showed less association with the parasite throughout the time course. Quantitative image analysis further supported these observations, showing significantly higher fluorescence intensity associated with *L. gasseri* compared to *G. vaginalis* (Fig. 7E). Collectively, these results demonstrate that *T. vaginalis* establishes a rapid, preferential, and time-dependent association with *L. gasseri* over *G. vaginalis*, underscoring the existence of selective parasite:bacteria interactions that extend beyond simple co-occurrence within the vaginal environment.

**Figure 7.**
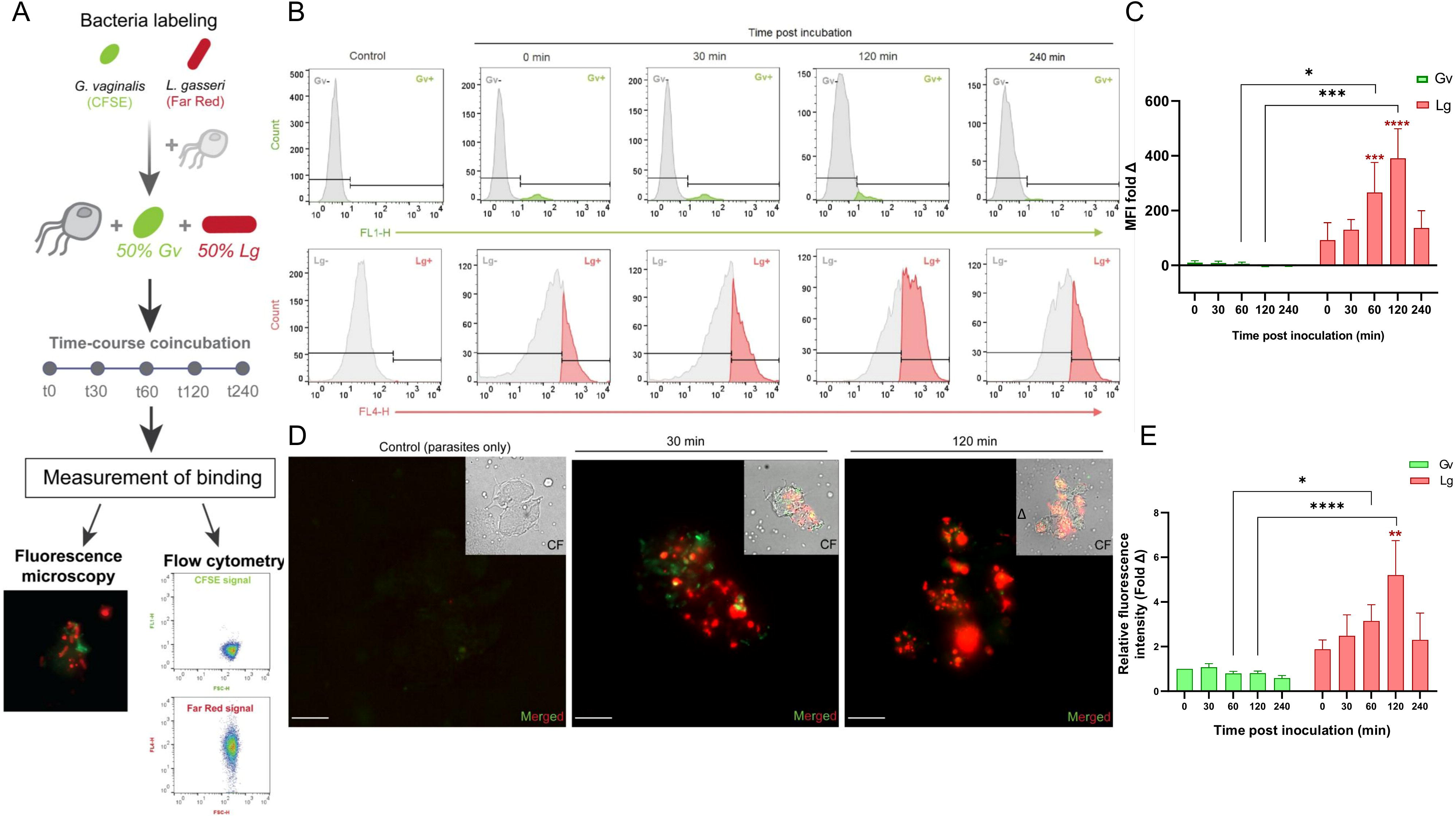
*T. vaginalis* exhibits preferential binding to *L. gasseri* over *G. vaginalis.* (A) Experimental scheme used to evaluate selective parasite: bacteria interaction. *Gardnerella vaginalis* (Gv) and *Lactobacillus gasseri* (Lg) were fluorescently labelled with CFSE (green) and CellTrace™ Far Red (red), respectively, mixed at a 1:1 ratio, and co-incubated with parasites for the indicated times (0–240 min). Then, bacterial binding to *T. vaginalis* was assessed by fluorescence microscopy and flow cytometry. Far red and CFSE signal from labelled *L. gasseri* and *G. vaginalis* respectively, analyzed by flow cytometry, is shown. (B) Representative flow cytometry histograms showing the association of fluorescently labeled bacteria with parasites at the indicated times post-incubation. Upper panels (FL1-H) correspond to the association of CFSE-labeled *G. vaginalis* (Gv+), and lower panels (FL4-H) correspond to the binding of Red–labeled *L. gasseri* (Lg+). Control histograms (parasites alone) are shown first, showing the background fluorescence (gray). (C) Median fluorescence intensity (Δmedian MFI) indicative of parasite-associated bacteria (Gv+, Lg+) at the indicated time points relative to time 0 for each bacterial species. Data represent mean ± SEM from independent experiments. Statistical significance is indicated (*P < 0.05; ***P < 0.001). (D) Representative fluorescence microscopy images of parasites incubated with labeled bacteria for 30 and 120 min. Merged images show the binding of Gv (green) and Lg (red) to the parasites. Insets show corresponding bright-field images. Scale bar, 10 µm. (E) Quantification of relative fluorescence intensity associated with parasites from microscopy images at the indicated time points relative to time 0 for each bacterial species. Data are shown as mean ± SD from of three independent biological experiments. Statistical analysis was performed using two-way ANOVA followed by multiple comparisons tests. Statistical significance is indicated as P < 0.05 (*), P < 0.001 (**), P < 0.0001 (****).

### *L. gasseri* induce *T. vaginalis* growth

We next asked whether these selective interactions and chemotaxis have functional consequences for parasite growth. *T. vaginalis* B7RC2 (Fig. 8D) and NYH209 (Fig. S1) strains were co-incubated with *L. gasseri, G. vaginalis,* or *E. coli*, and parasite growth was evaluated over a 24 h period. Parasite proliferation was influenced by the presence of *L. gasseri* but not by *G. vaginalis* or *E. coli*. Specifically, co-culture with *L. gasseri* resulted in a significant increase in *T. vaginalis* growth in B7RC2 (Fig. 8D) and NYH209 (Fig. S1) strains compared to control conditions. In contrast, co-incubation with *G. vaginalis* led to similar growth levels as control, whereas the presence of *E. coli* markedly impaired parasite proliferation. These results are consistent with previous findings describing inhibitory effects of *E. coli* on *T. vaginalis* growth (Chiu et al., 2025). These results indicate that the impact of bacterial co-culture on *T. vaginalis* growth is highly species specific.

**Figure 8.**
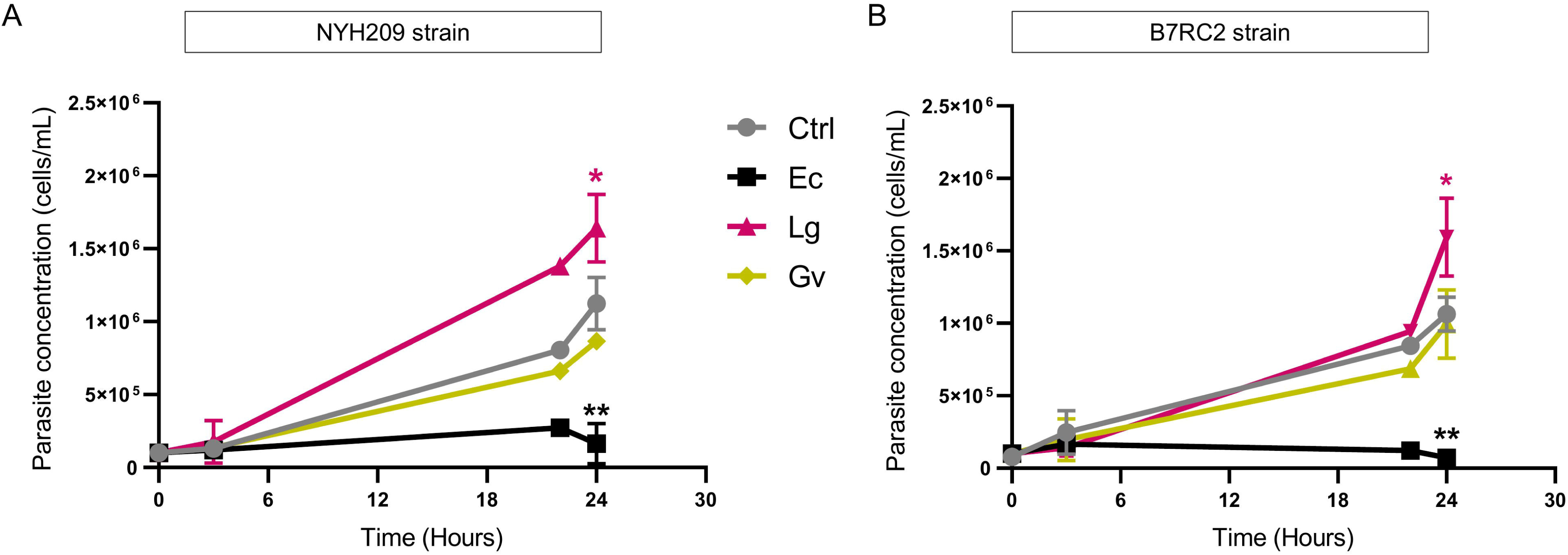
Effect of bacterial co-culture on *T. vaginalis* growth. Growth curves of the NYH209 (A) and B7RC2 (B) strains co-incubated with *L. gasseri* (Lg), *G. vaginalis* (Gv), or *E. coli* (Ec), compared to parasites growth in the absence of bacteria (Ctrl). Parasite concentration was measured over time and expressed as cell/mL. Statistical analysis was performed using mixed-effects models followed by Dunnett’s multiple comparisons test, comparing each bacterial condition to the control. Values are shown as mean ± SD from four biologically independent experiments. Statistical significance is indicated as P < 0.05 (*), P < 0.01 (**); ns, not significant.

## Discussion

In this study, we identify for the first time chemotaxis in *Trichomonas vaginalis* as a collective behavior that integrates environmental sensing with intercellular communication. The ability of the parasite to organize into coordinated multicellular assemblies and to migrate directionally in response to chemical gradients reveals a level of behavioral complexity not previously characterized in this extracellular pathogen. These findings position chemotaxis as a potentially important aspect of *T. vaginalis* persistence and pathogenicity. The coordinated migratory behavior observed in our assays is reminiscent of the phenomenon of social motility (SoMo) described in trypanosomes (Bastin & Rotureau, 2015; Oberholzer et al., 2010b; Shaw & Roditi, 2023). In SoMo, parasites do not move randomly; instead, groups of cells migrate collectively across semi-solid surfaces in highly organized projections, allowing the population to explore and colonize new environments (Bastin & Rotureau, 2015; Oberholzer et al., 2010b). In *Trypanosoma brucei*, SoMo is driven by the parasites’ ability to sense and respond to pH gradients that they generate themselves during growth. As trypanosomes metabolize nutrients, they acidify their local environment, creating a gradient across the surface of the plate (Shaw et al., 2022). Cells detect this gradient and migrate away from the acidic zone toward more alkaline regions, resulting in the characteristic outward radial movement observed in SoMo assays (Shaw et al., 2022). This behavior reflects a coordinated population level response in which parasite communities collectively interpret environmental cues to guide directional movement. Interestingly, our results indicate that, rather than migrating toward alkaline regions, *T. vaginalis* exhibits a strong attraction to acidic environments, which is consistent with the physiological conditions of the vagina in healthy women of reproductive age (pH 3.5 - 4.5) (Petrin et al., 1998; Vaneechoutte, 2017b). This fundamental difference suggests that *T. vaginalis* may be evolved with pH taxis as an adaptative ecological trait. The ability of *T. vaginalis* to sense and migrate toward acidic niches may therefore facilitate its vaginal colonization, promoting encounters with specific members of the microbiota and influencing host:bacteria:parasite interactions. Together, these observations suggest that chemotactic responses, analogous in concept to SoMo, may play an important role in shaping the ecological behavior of *T. vaginalis* within the vaginal mucosa. Although our results emphasize the role of pH as an important directional cue, it is highly probable that *T. vaginalis* chemotaxis likely integrates multiple physicochemical signals present in the vaginal environment.

A comprehensive understanding of host:parasite interactions, particularly in the case of extracellular pathogens such as *Trichomonas vaginalis*, requires considering not only the host environment but also the indigenous microbial communities present at the site of colonization. In the context of the human vagina, *T. vaginalis* does not exist in isolation but interacts with a diverse and dynamic microbial ecosystem, including commensal and potentially pathogenic bacteria. Our findings now reveal a selective chemotactic response toward both *Lactobacillus gasseri* and *Gardnerella vaginalis*, two bacterial species that dominate distinct vaginal community states. Although the parasite migrates toward both bacteria, the stronger attraction to *L. gasseri*, typically associated with a Lactobacillus-dominated eubiotic microbiota, is particularly intriguing. During eubiosis, lactobacilli acidify the vaginal microenvironment through high lactic acid production, and it has been proposed that this acidification contributes to pathogen exclusion, competitive adherence to epithelial cells, and inhibition of opportunistic microorganisms (Boris & Barbés, 2000; Tachedjian et al., 2017). In concordance with this proposal, it has been shown that the presence of Lactobacilli inhibited facultative bacteria, such as *G. vaginalis* and the anaerobes *P. anaerobius* and *P. bivia* (Atassi et al., 2006; Onderdonk et al., 2016; Phukan et al., 2018b; Skarin & Sylwan, 1986). Our findings suggest that *T. vaginalis* is capable of sensing this dominant local signal, i.e. lactic acid, as a cue to migrate toward lactobacilli-rich areas of the vaginal mucosa. Consistent with this idea of predatory-prey, the strong attraction to *L. gasseri* raises the possibility that *T. vaginalis* actively targets and kill protective lactobacilli, contributing to their depletion. Supporting this possibility, we showed that *T. vaginalis* exhibits a clear preferential binding to *L. gasseri* than *G. vaginalis* when the parasite is co-incubated with equal numbers of both bacteria. Therefore, this selective interaction could facilitate preferential predation on lactobacilli, potentially disrupting the protective vaginal microbiota. In concordance, previous studies have shown that *T. vaginalis* can ingest Lactobacillus species, including *L. gasseri (Juliano et al., 1991)*. This preferential binding also aligns with the observation that laterally acquired peptidoglycan hydrolases of the NlpC/P60 family were shown to empower the parasite to control lactobacilli (Pinheiro et al., 2018). This selective interaction might not be a stochastic event but could have evolved to facilitate a targeted predator-pray encounter specifically directed towards host-protective Lactobacilli.

This strategy of ‘targeted grazing’ finds parallels in other pathogens and phagocytes. For instance, *Entamoeba histolytica* utilizes its surface lectins to selectively bind and ingest specific bacterial populations, a process that significantly modulates its virulence and gut colonization (Ankri, 2021; Marie & Petri, 2014). Similarly, the environmental protist *Dictyostelium discoideum* exhibits a bacterial preference based on chemical sensing to ensure survival in complex microbial landscapes (Nasser et al., 2013; Rashidi & Ostrowski, 2019). Our findings suggest that *T. vaginalis* has evolved a mechanism that is selective for seeking and binding to lactobacilli allowing the parasite to shape the microbiome. This strategy aligns with clinical observations where *T. vaginalis* infection is strongly associated with the CST-IV microbiome, which lacks host-protective lactobacilli and resembles the dysbiotic polymicrobial community described for bacterial vaginosis (Brotman et al., 2012). While this link is well-documented, our data suggests that the shift toward dysbiosis is not just a prerequisite for infection, but a direct result of the parasite’s targeted chemotaxis and predation, helping explain this well-established link *(Amabebe & Anumba, 2018b; Dong et al., 2024; Z. (Sam) Ma & Li, 2017; Ravel et al., 2011)*. While BV has traditionally been viewed as a predisposing factor for *T. vaginalis* infection, our results now introduce a complementary explanation in which the parasite itself actively seek for host-protective lactobacilli in attempt to drive microbial destabilization. Through chemotactic migration and selective binding, *T. vaginalis* could target protective species, disrupt community structure, and promote CST-IV type dysbiosis. This mechanism aligns with earlier proposals of competitive exclusion between protozoa and lactobacilli (Barnett et al., 2023; Brotman et al., 2012; Matu et al., 2010; Pinheiro et al., 2018) and expands it by introducing chemotaxis and microbial sensing as critical active contributors. From this perspective, chemotaxis may function as an exploratory strategy that allows *T. vaginalis* to dynamically probe its microbial landscape and position itself within favourable microenvironments. Rather than responding passively to global vaginal conditions, the parasite appears capable of integrating localized physicochemical cues with the presence of specific bacterial populations. This behaviour may provide a selective advantage by coupling migration, resource access, and microbial competition at a microscale. The ability of *T. vaginalis* to recognize acidic gradients and selectively migrate toward specific bacterial species suggests that bacteria:parasite interactions are not merely competitive but may involve direct predation, altering community structure in ways that favor infection. Such interactions could destabilize vaginal homeostasis, promote dysbiosis, and influence disease severity, symptomatology, and transmission potential.

Previous studies have shown that lactobacilli can downmodulate *T. vaginalis* pathogenicity, suppress adherence, and limit parasite-induced inflammation (Fichorova et al., 2013; Phukan et al., 2018a) with *L. gasseri* ATCC 9857 as one of the most potent inhibitors of *T. vaginalis* adherence to epithelial cells. If these two microorganisms are natural competitors in the vagina, we now show that T. vaginalis is not passive, but this parasite hunts and binds to its pray with preference. This preference of bacterial encounter and predation could steadily create an inducive environment to *G. vaginalis* and other CST-IV bacteria species. This dysbiotic bacterial community has been repeatedly shown to enhance parasite pathogenesis (Artuyants et al., 2024; Fichorova et al., 2013; Hinderfeld et al., 2019). Prior research has demonstrated that *L. gasseri* and *G. vaginalis* exert opposing modulatory effects on *T. vaginalis* pathogenicity (Aquino et al., 2020; Fichorova et al., 2013, 2021; Hinderfeld et al., 2019; Phukan et al., 2013, 2018), further emphasizing the complex nature of parasite:microbiota interactions. By uncovering parasite-directed chemotaxis and selective interaction with vaginal bacteria, our study adds an additional layer to these observations. The interplay between parasite motility, physicochemical cues, and bacterial metabolites like lactic acid, likely shapes the spatial distribution of *T. vaginalis* within the vagina and influences infection dynamics.

Taken together, our findings position chemotaxis and selective binding as a central behavioral strategy enabling *T. vaginalis* to sense, navigate, and initiate downstream responses that could significantly reshape the vaginal environment including its microbial inhabitants. This work highlights the importance of considering vaginal microbiota as an integral element of host:parasite interactions. Further studies dissecting the molecular mechanisms underlying pH sensing, chemotaxis, and parasite:bacteria interactions will be essential for understanding the determinants of trichomoniasis and may identify new therapeutic strategies aimed at disrupting parasite:microbiota cross-talk and restoring vaginal homeostasis.

## Materials and Methods

### Parasites, bacteria, and culture media

*Trichomonas vaginalis* strains NYH209 (ATCC 50146) (Prat et al., 2024), B7RC2 (ATCC 50167; Greenville, NC, USA), CDC1132 (MSA1132; Mt. Dora, Fla, USA 2008) (Mercer et al., 2016) and B7268 (Upcroft & Upcroft, 2001) were cultured in Diamond’s trypticase-yeast extract-maltose medium supplemented with 10% heat-inactivated (HI) fetal bovine serum (FBS; NATOCOR, Argentina) and 10 U ml^−1^ penicillin and 10 μg ml^−1^ streptomycin (Gibco). Parasites were grown at 37°C and passaged daily. *Lactobacillus gasseri* ATCC 9857 and *Gardnerella vaginalis* ATCC 14018, kindly provided by Dr. Augusto Simões-Barbosa (University of Auckland, New Zealand), were cultured in modified Diamond’s medium (Hollander, 1976a)containing 0.1% Tween 80 (added prior to autoclaving) and supplemented after sterilization with 10% HI-FBS, at 37 °C under microaerophilic conditions. *Escherichia coli* strain OmniMAX (Invitrogen) was grown in Luria–Bertani (LB) medium with continuous agitation at 37 °C. Cultures were routinely subcultured daily to maintain log-phase growth.

### Social motility (SoMo) assay

Cultivation on semi-solid agarose plates was performed using adapted Diamond medium (Hollander, 1976b) supplemented with 10% HI-FBS and 1% antibiotics. Agarose was added directly to the medium at a final concentration of 0.54% (w/v), autoclaved, and allowed to cool to 37 °C for 1 h. A 20 ml aliquot of the agarose–medium solution was poured into Petri dishes, which were then air-dried without lids for 45 min in a laminar flow hood at room temperature. Parasites (3 × 10⁶ cells) were resuspended in 5 µl of medium and inoculated onto the surface of the agarose plates. Plates were subsequently incubated in an anaerobic chamber at 37 °C and monitored daily until colonies became visible. Differences in parasite distribution on the plates reflected the experimental design and inoculation layout, while the number of parasites inoculated was kept constant across all conditions.

### pH taxis and chemotactic assays on semi-solid agarose plates

Parasites were inoculated onto semi-solid agarose plates as previously described for social motility assays. Chemical stimuli were applied to the agarose surface at a distance of 2 cm from the central site of parasite inoculation. Plates were incubated at 37°C under anaerobic conditions and monitored daily until directed parasite migration toward specific pH conditions became apparent.

Acidic microenvironments were generated using HCl, lactic acid, or acetic acid, whereas alkaline conditions were established using NaOH. All chemical stimuli were prepared from 1 M stock solutions, diluted in sterile water, and freshly prepared prior to each experiment. Parasite migration toward or away from the applied stimuli (10 µl of each solution) was documented by imaging. The spatial distribution of chemical solutions on the agarose surface was adjusted according to the experimental design, with stimuli positioned either at opposing locations or in a triangular configuration, as indicated in the corresponding figures.

### Bacterial preparation and inoculation

Bacterial cultures were grown to logarithmic phase and the optical density at 600 nm (OD_600_) was measured immediately prior to each experiment. To ensure comparable bacterial inputs across conditions, the inoculum were normalized based on OD_600_ using species-specific conversion factors (cells per mL per OD unit): *Escherichia coli*, 8 × 10⁸ cells mL^-^¹ OD^-^¹; *Lactobacillus gasseri* and *Bacillus sp.* (MT3), 1 × 10⁹ cells mL^-^¹ OD^-^¹; and *Gardnerella vaginalis*, 5 × 10⁸ cells mL^-^¹ OD^-^¹, as previously described (Aponte et al., 2024; X. Ma et al., 2022; Mira et al., 2022). For each bacterial species, the volume corresponding to 4 × 10⁸ cells was centrifuged at 4000 × g for 10 min, washed once in fresh medium and resuspended in 10 µl prior to inoculation. For migration assays, NYH209 or B7RC2 strains were inoculated at the centre of semi-solid agarose plates using the same conditions and parasite density described for the SoMo assay, except that antibiotics were omitted, and bacterial suspensions were placed at fixed peripheral positions arranged in a triangular configuration. Plates were incubated under anaerobic conditions and monitored over time until directional sensing by the parasite toward bacterial-derived cues became evident.

### Bacterial growth and medium acidification assay

To evaluate medium acidification associated with bacterial growth, *Escherichia coli*, *Gardnerella vaginalis* and *Lactobacillus gasseri* were cultured in modified Diamond medium supplemented with 0.1% Tween 80 and 10% FBS, as described above. Overnight bacterial cultures were adjusted to an initial optical density of OD_600_ = 0.2. *L. gasseri* and *G. vaginalis cultures* were incubated at 37 °C under microaerophilic conditions without agitation, whereas *E. coli* cultures were incubated at 37 °C with agitation. The initial pH of the culture medium was 6.2, corresponding to Diamond medium. At defined time points, aliquots of the cultures were collected, and the pH of the supernatants was measured using a calibrated pH electrode. Medium-only controls were incubated under the same conditions and processed in parallel. Changes in pH over time were used as a readout of bacterial-mediated acidification of the culture medium.

### Parasite growth during bacteria co-culture assays

To assess interactions between *Trichomonas vaginalis* and vaginal bacteria, co-culture assays were performed using strains NYH209 or B7RC2 and the bacterial species *Lactobacillus gasseri, Gardnerella vaginalis, or Escherichia coli*. Parasites were seeded at a density of 1 × 10⁵ cells/ml, while bacterial cultures were adjusted to an initial optical density of OD_600_ = 0.2. Co-cultures were established in Diamond medium supplemented with 10% HI-FBS. Parasite–bacteria mixtures were incubated at 37°C under microaerophilic conditions without agitation. Control cultures containing parasites or bacteria alone were included in parallel under identical conditions. At the indicated time points, parasite density was quantified by cell counting using a CellDrop™ automated cell counter (DeNovix).

### Parasite-bacteria interaction by Flow Cytometry

Association between *T. vaginalis* and vaginal bacteria was analyzed by flow cytometry using differentially fluorescently labelled bacteria. *L. gasseri* and *G. vaginalis* were grown to logarithmic phase and normalized to an initial optical density (OD_600_) of 0.2. Bacterial cells were collected by centrifugation at 4,000 × g and resuspended in 1× PBS for fluorescent labelling. *L. gasseri* was stained with CellTrace Far Red (Invitrogen; 1 µM final concentration) for 20 min at room temperature with gentle agitation, whereas *G. vaginalis* was labeled with CFSE (Sigma; 40 µM final concentration) for 20 min at 37 °C. After staining, bacteria were washed 3–5 times with PBS to remove excess dye. Fluorescent labelling efficiency and relative bacterial abundance were verified by flow cytometry prior to co-incubation. For parasite–bacteria interaction assays, equal proportions of fluorescently labelled *L. gasseri* and *G. vaginalis* were mixed (50:50) and co-incubated with non-labeled *T. vaginalis* (5 × 10⁵ parasites/ml) in Diamond medium without serum supplementation. Co-incubations were performed for the indicated time points before flow cytometry analysis. To control for background fluorescence and dye carryover, control samples containing PBS incubated with each dye in the absence of bacteria were included. Flow cytometry acquisition was performed by detecting CFSE and CellTracker™ Far Red fluorescence in the FL1 and FL4 channels, respectively. Data were analysed by calculating changes in median fluorescence intensity (Δmedian MFI) associated with parasite populations.

### Fluorescence microscopy of parasite–bacteria association

*L. gasseri* and *G. vaginalis* were fluorescently labeled with CellTracker Far Red and CFSE respectively, and co-incubated with non-labeled *T. vaginalis* (5 × 10⁵ parasites/ml) in Diamond medium without serum supplementation for 4 h, as described above. At different time points, samples were collected by centrifugation at 4,000 × g for 10 min and fixed with 4% paraformaldehyde. Fixed samples were mounted using Fluoromount™ and examined by fluorescence microscopy using a Zeiss Axio Observer 7 inverted microscope (Zeiss). Images were acquired using identical acquisition settings across conditions. Microscopy experiments were performed in three independent biological replicates.

### Statistical analysis

Statistical analyses were performed using GraphPad Prism for Windows (version 8.0.2). The choice of statistical test depended on the experimental design and data structure of each assay and is specified in the corresponding figure legends. One-way or two-way analysis of variance (ANOVA), as appropriate, was used to evaluate differences in parasite migration in response to bacterial stimuli and pH-related experiments. When significant effects were detected, appropriate post hoc multiple comparison tests were applied. Linear mixed-effects models (REML) were used for datasets involving repeated measurements over time or multiple sources of experimental variability, including parasite density measurements, flow cytometry analyses, and fluorescence microscopy quantifications, with experimental replicates treated as random effects when applicable. Data are presented as mean ± standard deviation (SD). Statistical significance was established at P < 0.05.

## Acknowledgments

The authors thank Jose Luis Burgos and Carla Chrestia for his technical assistance and our colleagues in the lab for helpful discussions. This research was supported with a grant from ANPCyT grant BID PICT-2019-01671 (to N.d.M.). N.d.M. is a researcher from the National Council of Research (CONICET) and the National University of San Martin (UNSAM). M.B.P. is a PhD fellow from CONICET. Research was also supported by a Prebys Research Heroes Award from the Prebys Foundation to A.M.R and a Research Project Grant to A.M.R from the National Institute on Minority Health and Health Disparities of the National Institutes of Health under Award Numbers U54MD012397 (SDSU HealthLINK Center) and S21MD010690 (SDSU HealthLINK Endowment). The funders had no role in study design, data collection and analysis, decision to publish, or preparation of the manuscript.

